# Genome-wide characterization of *Salmonella* Typhimurium genes required for the fitness under iron restriction

**DOI:** 10.1101/2021.07.28.454258

**Authors:** Sardar Karash, Tieshan Jiang, Young Min Kwon

## Abstract

Iron is a crucial element for bacterial survival and virulence. During *Salmonella* infection, the host utilizes a variety of mechanisms to starve the pathogen from iron. However, *Salmonella* activates distinctive defense mechanisms to acquire iron and survive in iron-restricted host environments. Yet, the comprehensive set of the conditionally essential genes that underpin *Salmonella* survival under iron-restricted niches have not been fully explored. Here, we employed transposon sequencing (Tn-seq) method for high-resolution elucidation of the genes in *Salmonella* Typhimurium (*S.* Typhimurium) 14028s strain required for the growth under the *in vitro* conditions with 4 different levels of iron restriction achieved by iron chelator 2,2′-Dipyridine (Dip): mild (100 and 250), moderate (250) and severe iron restriction (400 μM). We found that the fitness of the mutants reduced significantly for 28 genes, suggesting the importance of these genes for the growth under iron restriction. These genes include *sufABCDSE*, iron transport *fepD*, siderophore *tonB*, sigma factor E *ropE*, phosphate transport *pstAB,* and zinc exporter *zntA*. The siderophore gene *tonB* was required in mild and moderate iron-restricted conditions, but they became dispensable in severe iron-restricted conditions. Remarkably, *rpoE* was required in moderate and severe iron restrictions, leading to complete attenuation of the mutant under these conditions. We also identified 30 genes for which the deletion of the genes resulted in increased fitness under iron-restricted conditions. The findings broaden our knowledge of how *S*. Typhimurium survives in iron-deficient environments, which could be utilized for the development of new therapeutic strategies targeting the pathways vital for iron metabolism, trafficking, and scavenging.

## Introduction

Iron is a cornerstone for numerous cellular metabolisms and serves as a cofactor for some proteins with vital functions. Iron is involved in many critical biochemical reactions, including respiration, tricarboxylic acid cycle, synthesis of metabolites, and enzyme catalysis. Therefore, iron is a crucial metal for the survival of bacterial pathogens [1, 2]. The non-typhoidal intracellular *S.* Typhimurium can infect a wide range of hosts and cause gastroenteritis [3]. It has been estimated that *S*. Typhimurium is accountable for 93.8 million cases of gastroenteritis, leading to 155,000 deaths worldwide yearly [4]. As iron accessibility is vital for *S*. Typhimurium pathogenesis, the host uses a variety of mechanisms to sequester it from bacteria [5]. Also, it has been shown that a probiotic *Escherichia* coli Nissle 1917 reduces *S.* Typhimurium colonization by competing for iron [6]. After consuming foods or water contaminated with *Salmonella*, the pathogen reaches the intestine and breaches epithelial tissue to enter macrophages [7]. A defense mechanism that the host uses to fight against pathogens is depleting free iron via iron-sequestering proteins such as heme, hepcidin, ferritin, transferrin, and lactoferrin [8]. Hepcidin is produced in response to infection to decrease the concentration of iron in the plasma and facilitate sequestration of iron in macrophages [2]. Despite a widespread counter-defensive strategy of hosts against the pathogens, *S.* Typhimurium thrives in the inflamed gut and can survive and replicate in macrophages [6, 7]. Bacterial pathogens, including *Salmonella*, employ aggressive acquisition processes to scavenge iron from the hosts through the synthesis and excretion of high-affinity iron chelators named siderophores [9]. It has been also suggested that modulating host iron homeostasis may be a path to tackle multidrug-resistant intracellular bacteria [10]. Still, our understanding of the genes in *S.* Typhimurium that are required for survival in iron-restricted environments is incomplete.

It is highly important to characterize the entire genome of *S.* Typhimurium in a biologically relevant range of iron restriction to gain a comprehensive understanding of the genes and their proteins that play a role in coping with the stressor. The 2,2‵-Dipyridyl (Dip) is the most commonly used, membrane-permeable iron chelator and selective agent to chelate Fe^2+^ [11]. In a previous study, the promoters in *S*. Typhimurium that respond to 200 µM Dip were identified using a high-throughput approach based on the random promoter fusions [12]. Microarray also has been used intensively in different bacteria to profile global transcriptional responses to iron limitation, using varying concentrations of Dip: for instance, 200 µM (*E*. *coli*) [13], 160 µM (*Shewanella oneidensis*) [14]; 300 µM (*Actinobacillus pleuropneumoniae*) [15]; 40 µM (*Leptospira interrogans*) [16]; 200 µM (*Acinetobacter baumannii*) [17] and 200 µM (*S*. Typhimurium) [18]. RNA-seq has also been applied for transcriptomic responses to 30 µM Dip for *Rhodobacter sphaeroides* [19] and 200 µM Dip for *S*. Typhimurium [20]. In these studies, the tested bacteria were typically exposed to one selected concentration of Dip for a short time to explore the gene expression responses. On the contrary, in our current study, we allowed the genome-saturating Tn5 mutant libraries of *S*. Typhimurium to grow in the presence of varying levels of iron-restricted conditions and quantitatively tracked all mutants under these stress conditions to gain more direct evidence for the genes that are required for the fitness under these stress conditions.

Utilizing highly saturated Tn5 libraries and more than a quarter-billion reads from Tn5-genomic junctions, we identified the conditionally essential genes in *S*. Typhimurium that are required for the growth under varying levels of iron restriction. We demonstrated that *sufABCDSE* operon is important for bacterial fitness under moderate (250 µM) and severe (400 µM), but not under mild iron restriction conditions (100 and 150 µM Dip). We also found new genes that are critical for the growth under iron-restricted conditions, including the genes encoding sigma factor E and the proteins in electron transport, glycolysis and gluconeogenesis, phosphate transport, and zinc export. Finally, we also identified the genes that when deleted increase the mutant fitness under iron restriction. The genes identified in this study can be exploited as targets for the development of novel antibiotics and expand our knowledge related to iron acquisition and trafficking in *S*. Typhimurium.

## Results and Discussion

### *S.* Typhimurium growth response to different concentrations of 2,2‵-Dipyridyl

Initially, we investigated the growth response of the wild-type *S*. Typhimurium 14028s to different concentrations of iron chelator Dip. The examined Dip concentrations ranged from 100 to 2,000 μM. As illustrated in **Fig. S1**, the final optical density (OD_600_) of the bacterial cultures after 18 hr incubation at 37°C reduced as the concentration of Dip increased. The bacteria did grow in the presence of Dip at the concentrations of 100 to 500 μM. But at 1,000 μM Dip and above the bacteria could hardly grow with only a marginal increase in the optical density at 1,000 μM. We found a significant decrease of OD_600_ in the presence of 100 μM Dip as compared to the control culture with no Dip (*p* < 0.05). This was an indicator that the 100 μM Dip had a negative effect on the growth of *S*. Typhimurium. Therefore, we decided to use Dip concentrations ranging from 100 to 400 μM for the following Tn-seq selections; the concentrations of 100, 150, 250, and 400 μM chosen for selections of the Tn5 library are hereafter referred to Dip100, Dip150, Dip250 (Dip250-I and Dip250-II for 2 independent Tn-seq selections), and Dip400, respectively (**Fig. S2**). As the concentration of Dip increased, the growth rate reduced, and maximum OD_600_ decreased (**Fig. 1A** and **Fig. 1B**, respectively). The Dip250 had a profound effect on the growth and it reduced maximum OD_600_ by 34% in reference to LB control (**Table S1**). Dip400 had a severe effect on *S*. Typhimurium growth; it reduced the growth rate by 26% and maximum OD_600_ by 48%. Since *Salmonella* encounters host niches with different concentrations of available iron, we reasoned that our Tn-seq selection conditions representing a wide range of iron restriction are more relevant in revealing the strategies *Salmonella* employs to cope with iron-restriction stress during infection in the host as compared to one condition with a fixed level of iron restriction.

**Figure 1.**
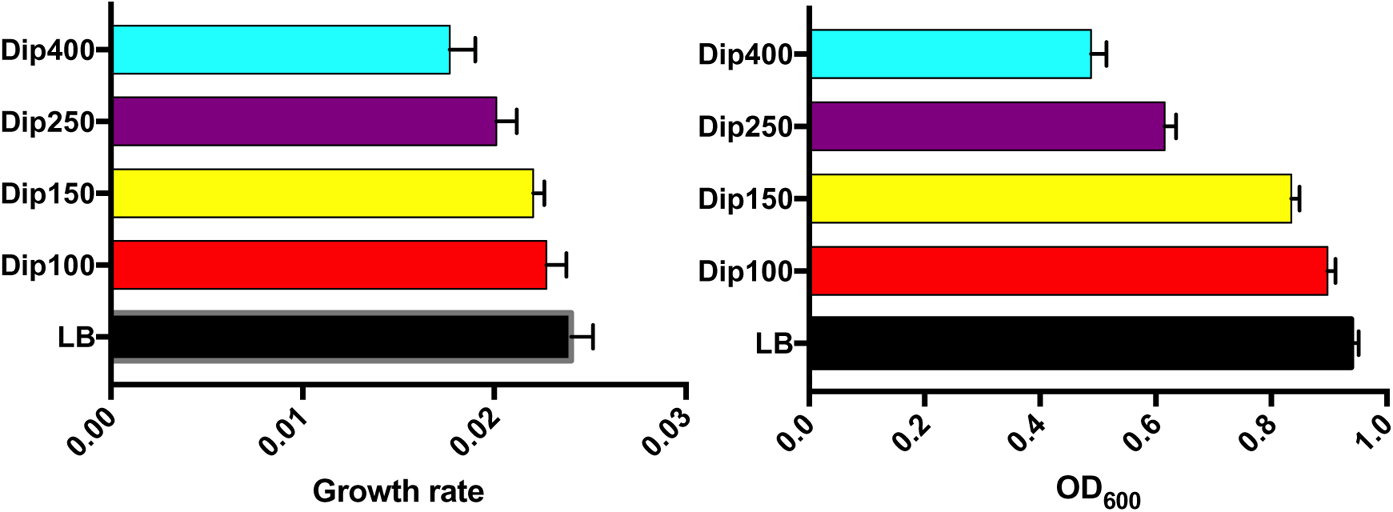
Effect of varying concentrations of 2,2‵-Dipyridyl on the growth of *S*. Typhimurium 14028s. *S*. Typhimurium 14028s wild-type strain was grown in LB broth supplemented with different levels of Dip (0, 100, 125, 250, or 400 µM). The cultures were incubated in a 96-well plate and OD_600_ was measured with Tecan Infinite 200 microplate reader for 24 hr at 37°C. The collected data were used to calculate the growth rate and to obtain a maximum OD_600._ Data represent at least three replicates.

**Figure 2.**
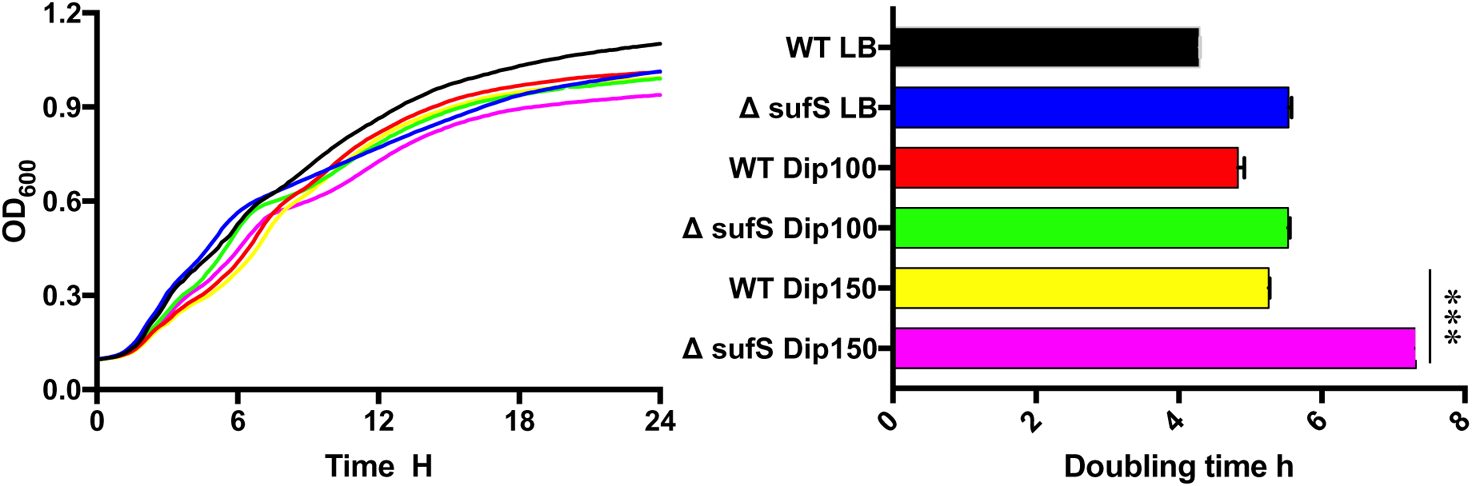
*sufS* is required for the optimal growth of *S*. Typhimurium under iron-restricted conditions. *S*. Typhimurium 14028s wild-type and Δ*sufS* mutant were grown in LB broth, supplemented with 0 (control), 100 or 150 µM 2,2‵-Dipyridyl (Dip). OD_600_ was recorded every 15 minutes during incubation at 37°C for 24 hr in a 96-well plate. Data represent at least three replicates. Statistical significance of doubling times was determined by unpaired two-tailed *t* test, \*\*\**p* < 0.001.

**Table 1.** The genes in S. Typhimurium 14028s that are required for fitness under iron restriction conditions

| Genes<br>(with Tn5<br>insertions) | Fitness (log<sub>2</sub>FC) |  |  |  | Previously implicated role in iron-restriction |  |  |
| --- | --- | --- | --- | --- | --- | --- | --- |
|  | Dip100 | Dip150 | Dip250 | Dip400 | *S. Typhimurium*<br>(RNA-seq)<sup>20</sup> | *S. Typhimurium* | *E. coli* |
| *degS* | -0.08 | -0.75 | -2.60 | -8.35 | No | No | No |
| *exbB* | -1.4 | -3.28 | -1.80 | -1.81 | Yes | No | Yes<sup>45</sup> |
| *fepD* | -2.91 | -4.27 | -3.56 | -0.22 | No | Yes<sup>44</sup> | Yes<sup>43</sup> |
| *fre* | -0.97 | -1.04 | -1.50 | 0.03 | No | No | Yes<sup>37</sup> |
| *ggt* | -0.97 | -1.11 | -0.66 | -1.24 | No | No | No |
| *gpmA* | 0 | 0 | -3.09 | +0.36 | No | No | No |
| *hemL* | -4.52 | -4.52 | -2.55 | -0.24 | No | No | No |
| *ndh* | -0.07 | -1.62 | -1.91 | -1.10 | No | No | No |
| *osmE* | -0.38 | -0.25 | -1.00 | -1.35 | No | No | No |
| *pstA* | -0.56 | -0.45 | -1.45 | -0.57 | No | No | No |
| *pstB* | -0.77 | -1.10 | -1.39 | -2.04 | No | No | No |
| *rpoE* | -0.43 | -1.53 | -5.96 | -7.24 | No | No | No |
| *sodA* | -1.40 | -1.17 | -2.59 | -1.24 | No | Yes<sup>46</sup> | Yes<sup>56</sup> |
| *STM14_4330* | -0.04 | -0.19 | -0.40 | -0.63 | No | No | No |
| *STM14_4612* | -0.09 | +0.11 | -0.49 | -0.45 | - | No | - |
| *sufA* | -0.03 | -0.39 | -1.46 | -2.49 | Yes | No | Yes<sup>23</sup> |
| *sufB* | +0.07 | +0.28 | -1.67 | -2.84 | Yes | No | Yes<sup>23</sup> |
| *sufC* | -0.11 | -0.16 | -1.60 | -2.67 | Yes | No | Yes<sup>23</sup> |
| *sufD* | +0.13 | -0.22 | -2.17 | -2.75 | Yes | No | Yes<sup>23</sup> |
| *sufS* | -0.46 | -0.24 | -1.91 | -2.63 | Yes | No | Yes<sup>23</sup> |
| *tonB* | -0.58 | -6.43 | -4.31 | +1.44 | No | Yes<sup>46</sup> | Yes<sup>45</sup> |
| *ydgM (rsxB)* | -0.33 | -1.16 | -0.94 | +0.05 | No | No | No |
| *yfhP (iscR)* | 3.09 | -1.58 | -2.75 | -2.75 | Yes | Yes<sup>27</sup> | Yes<sup>26</sup> |
| *ygjQ* | -0.33 | -0.56 | -1.01 | -1.19 | No | No | No |
| *yhfK* | 0 | +0.72 | -0.61 | -0.44 | No | No | No |
| *yhgI (nfuA)* | -0.39 | -0.15 | -1.91 | +0.53 | No | No | Yes<sup>32</sup> |
| *ynhA (sufE)* | +0.27 | -0.02 | -1.91 | -2.16 | Yes | No | Yes<sup>23</sup> |
| *zntA* | -0.70 | -1.39 | -0.63 | -1.03 | No | No | No |

Dip represents 2,2'-Dipyridyl and the numbers indicate the concentrations of Dip (μM). The mutants with significantly reduced fitness are shaded with green color ( $p < 0.05$ ). In the RNA-seq study (reference 20), *S. Typhimurium* was exposed to 200 μM Dip for 10 min in Lennox broth.

### Selection conditions and summary of Tn-seq reads

Initially, we constructed a transposon Tn5 mutant library which consists of 325,000 random mutants as described previously [21]. Briefly, these transposon mutants were recovered on 50 LB agar plates (Library-A). After sequencing, it turned out that there was an insertion in every 42 nucleotides on average in the *S*. Typhimurium chromosome and 90% of ORFs had insertions. Then, we made another library containing 325,000 Tn5 mutants (Library-B) and combined it with Library-A, forming Library-AB. As a result, there was an insertion per 25 nucleotides on average and 92.6% of ORFs had insertions in Library-AB. The study design for the mutant selections using both Library-A and Library-AB is illustrated in **Fig. S2**. The respective controls were LB-II, and LB-III and the varying levels of iron-limited conditions were Dip100, Dip150, Dip250-I, Dip250-II, and Dip400 (**Fig. S2**). For each of the mutant selections, 20 ml of LB broth in a 300 ml flask containing an appropriate concentration of Dip was inoculated with the relevant transposon library. The seeding CFUs were approximately 10 cells for each mutant. Then, the cultures were grown until the culture reached the mid-log phase. LB-II and LB-III required about 5 hr to reach the mid-exponential phase, while this time was extended as the concentration of Dip increased. It took 10 hr and 12 hr for Dip250 and Dip400, respectively (**Table S2**). Next, the genomic DNA was extracted from each culture, and Tn-seq amplicon libraries were prepared for HiSeq sequencing as described previously [22].

**Table 2.** The genes in S. Typhimurium 14028s that upon deletion increased the fitness of the mutants under iron restriction conditions

| Genes<br>(with Tn5<br>insertions) | Biological Functions | Fitness (log<sub>2</sub>FC) |  |  |  |  |
| --- | --- | --- | --- | --- | --- | --- |
|  |  | Dip100 | Dip150 | Dip250-I | Dip250-II | Dip400 |
| *acnB* | Tricarboxylic acid cycle | 0.68 | 0.75 | 0.83 | 0.77 | 1.34 |
| *atpA* | ATP synthesis | 0.67 | 0.37 | 5.05 | 4.99 | 4.86 |
| *eda* | Carbohydrate metabolism | 1.00 | 1.00 | 3.03 | 3.75 | 3.62 |
| *fabF* | Fatty acid biosynthesis | 0.37 | 0.49 | 1.52 | 2.24 | 1.49 |
| *guaB* | Purine biosynthesis | 0.64 | 0.58 | 1.59 | 1.29 | 1.28 |
| *htpX* | Membrane-localized protease | 0.97 | 0.97 | 1.79 | 1.02 | 1.82 |
| *icdA* | Tricarboxylic acid cycle | 0.90 | 0.92 | 4.19 | 3.54 | 4.64 |
| *nlpB* | Outer membrane protein assembly factor | 0.69 | 0.93 | 0.7 | 1.1 | 1.75 |
| *ompW* | Transmembrane transport | 0.75 | 0.55 | 1.44 | 1.02 | 0.71 |
| *pnp* | Deoxyguanosine catabolic process | 0.94 | 0.88 | 2.49 | 2.7 | 1.49 |
| *purA* | Purine nucleotide biosynthetic process | 0.69 | 0.42 | 1.14 | 1.05 | 1.95 |
| *rfaC* | Lipopolysaccharide biosynthesis | 0.13 | 0.20 | 0.89 | 3.13 | 0.39 |
| *rfbB* | Lipopolysaccharide biosynthesis | 0.62 | 0.86 | 0.65 | 0.86 | 0.69 |
| *rfbH* | Lipopolysaccharide biosynthesis | 0.46 | 0.82 | 0.63 | 0.46 | 1.15 |
| *rfbI* | Lipopolysaccharide biosynthesis | 0.97 | 0.89 | 0.83 | 0.97 | 0.69 |
| *rprA* | Putative regulator | 0.12 | 0.10 | 6.49 | 0.06 | 0.33 |
| *sapA* | Peptide transmembrane transporter | 0.92 | 0.52 | 0.66 | 0.59 | 1.32 |
| *sdhD* | Tricarboxylic acid cycle | 0.50 | 1.00 | 1.94 | 1.09 | 2.16 |
| *smvA* | Transmembrane transport | 0.50 | 0.35 | 1.46 | 1.31 | 0.96 |
| *STM14_0726* | Integral component of membrane | 0.33 | 0.66 | 1.28 | 1.81 | 3.9 |
| *STM14_1429* | Putative regulator | 0.33 | 0.10 | 4.51 | 1.99 | 5.99 |
| *STM14_2094* | Putative catalase | 0.63 | 0.52 | 1.26 | 1.55 | 1.31 |
| *STM14_2709* | Gluconeogenesis | 0.85 | 0.97 | 0.82 | 0.95 | 1.11 |
| *trpE* | Amino-acid biosynthesis | 0.60 | 0.19 | 0.96 | 1.14 | 1.05 |
| *tuf_1* | Protein biosynthesis | 0.68 | 0.70 | 2.21 | 0.33 | 2.95 |
| *umuD* | DNA repair | 0.59 | 0.85 | 2.07 | 0.5 | 2.05 |
| *yaeD* | Carbohydrate metabolic process | 0.33 | 1.00 | 5.07 | 4.33 | 2.44 |
| *ycfC* | High frequency lysogenization protein | 0.66 | 0.70 | 3.4 | 3.17 | 1.73 |
| *ychH* | Integral component of membrane | 1.00 | 1.00 | 1.63 | 2.49 | 0.27 |
| *yeiB* | Integral component of membrane | 0.87 | 0.90 | 1.11 | 1.38 | 0.74 |

Dip represents 2,2'-Dipyridyl and the numbers indicate the concentrations of Dip (μM). The mutants with significantly increased fitness are shaded with green color ( $p < 0.05$ ).

In this study, we successfully mapped more than 173 million sequence reads to the chromosome of *S*. Typhimurium 14028s (NC_016856.1) for all selected conditions combined. The mean length of mapped genomic junction sequences was 91 nucleotides long. The highly saturated mutant library (Library-AB) contained 193,728 unique insertions (**Table S3**). The high number and long length of the mapped reads allowed us to identify conditionally essential genes with high precision. Previously, we showed that our Tn-seq protocol is highly reproducible [21]. In this study, a comparison of the Tn-seq profiles between the two biological replicates, Dip250-I and Dip250-II, indicated that the correlation coefficient (*r*) of unique insertions per ORF between was 0.995 (**Fig. S3A**). In addition, the *r* of essentiality indices per ORF as calculated by Tn-seq Explorer [23] between Dip250-I and Dip250-II was 0.990 (**Fig. S3B**). These results indicate the robustness and reproducibility of our Tn-seq library protocol used in this study.

### Genes implicated in the growth under iron-restricted conditions

Combining the results from all Tn-seq analyses under varying levels of iron-restricted conditions, we identified 58 genes for which the deletion mutants displayed either increased or reduced fitness in response to different concentrations of iron chelator Dip. Mutant fitness was reduced for 28 genes, while increased for 30 genes (**Supplementary Data set 1**). In other words, the 28 conditionally essential genes are required for the robust growth of *S*. Typhimurium under iron-restricted conditions. On the contrary, the 30 genes exhibited increased mutant fitness when deleted. We also identified essential genes of *S*. Typhimurium under these conditions, and the details of the findings were reported previously [24].

### Conditionally essential genes required for fitness under iron-restricted conditions

Our results of genome-wide analyses using Tn-seq show that 28 conditionally essential genes of *S*. Typhimurium are required for growth under different levels of iron restriction. Some of the genes have been implicated to have a role in growth under iron-restricted conditions in previous studies, but other genes have never been associated with iron restriction (**Table 1**).

#### Iron-sulfur cluster assembly genes

We found that iron sulfur cluster assembly operon *sufABCDSE* is required for the growth of *S*. Typhimurium under iron-restricted conditions (**Table 1**). The fitness of all six genes in *suf* operon was reduced significantly. Although the genes in *suf* operon are required for the bacteria to combat the iron-deficient environments, the fact that numerous reads corresponding to the Tn5 insertions in these genes were detected indicates these genes are not essential (**Supplementary Data set 2**). To confirm this finding, we examined the growth of *S*. Typhimurium lacking *sufS* under iron-restricted conditions (Dip100 and Dip150). The result indicated that the doubling time of Δ*sufS* deletion mutant were significantly higher as compared to the wild-type under Dip150 (*p* < 0.001)(**Fig. 2**). *E*. *coli* and *Salmonella* have two Fe-S cluster assembly systems, *isc,* and *suf*. Under iron-restricted conditions, *E*. *coli* utilizes *suf* operon for Fe–S cluster assembly [25]. *sufC* of *Salmonella* Typhi was previously shown to be required for survival in macrophages [26]. The ortholog of *sufS* in *Mycobacterium tuberculosis* is implicated in iron metabolism because Δ*sufS* mutant of *M*. *tuberculosis* exhibited longer doubling time in the presence of Dip [27]. In addition to the vital role of *suf* system, our Tn-seq data also indicated that *iscR* is required for *S*. Typhimurium growth under iron-restricted conditions (Table 1). The *suf* operon is controlled by *iscR* in *E*. *coli* [28]. IscR is not only involved in Fe-S cluster biogenesis but also implicated as a pleiotropic transcriptional regulator. *S*. Typhimurium *iscR* regulates SPI-I TTSS genes (*Salmonella* pathogenicity island 1 type III secretion system) when the iron level is low *in vivo* [29]. *Pseudomonas aeruginosa* IscR is considered a global regulator and it senses Fe-S cluster proteins. *P*. *aeruginosa* possesses only iron-sulfur cluster (ISC) system [30] and Δ*iscR* mutant showed attenuated virulence in *Drosophila melanogaster* and mouse peritonitis models [31]. Further, it was found that IscR regulates the expression of more than 40 genes involved in Fe-S cluster homeostasis in *E*. *coli* [32]. Therefore, we speculate that IscR is required for fitness in *S.* Typhimurium through its regulation of *suf* operon. We conducted protein-protein interaction network analysis and the result also indicates that four Suf proteins (SufBCDS) interact with IscR (**Fig. S4**). The molecular mechanisms of Fe-S proteins in bacteria have been extensively reviewed elsewhere [32]. Moreover, *nfuA* (*yhgI*) encodes an Fe-S cluster carrier protein, which was reported as a scaffold/chaperone for damaged Fe-S cluster proteins in *E*. *coli* [33]. Our Tn-seq screening shows that *nfuA* is required for *S*. Typhimurium growth under the iron-restricted condition. The Δ*nfuA* mutant in *P*. *aeruginosa* was sensitive to Dip and less virulent in *C*. *elegans* [34]. Deletion of *nfuA* in *E*. *coli* also caused the mutant to be susceptible to iron depletion [35]. The ortholog of *nfuA* in *Acinetobacter baumannii* plays a role in intracellular iron hemostasis and the bacterium cannot grow in iron-chelated media when the gene is deleted [36]. Here we demonstrate for the first time that *nfuA* is also critical for *S*. Typhimurium growth under iron-restricted conditions, and we speculate the protein is possibly involved in Fe-S cluster biogenesis.

**Figure 4.**
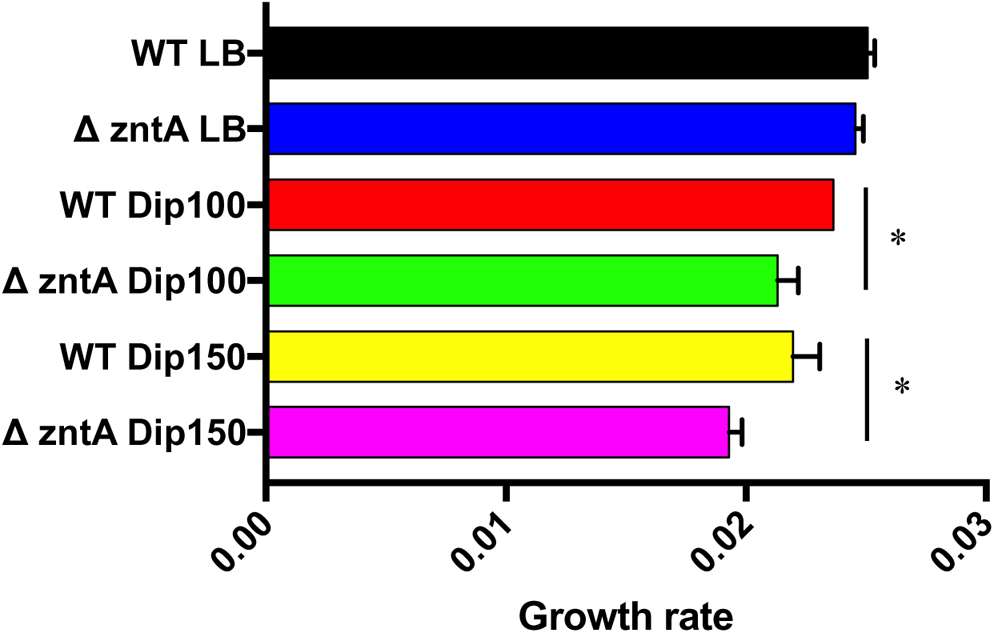
*zntA* is required for the growth of *S*. Typhimurium under iron-restricted conditions. *S*. Typhimurium 14028s wild-type and Δ*zntA* mutant were grown in LB broth supplemented with 0 (control), 100 or 150 µM 2,2‵-Dipyridyl (Dip). OD_600_ was recorded every 15 minutes during incubation at 37°C for 24 h in a 96-well plate. Statistical significance was determined by unpaired two-tailed *t* test, \**P* < 0.05.

#### Iron homeostasis

NAD(P)H-flavin reductase, *fre*, was also required under iron-restricted conditions (**Table 1**). Fre protein likely reduces free flavins, and consequently, the lower level of flavins reduces ferric iron to ferrous iron in *E*. *coli* [37, 38]. We speculate that the protein product of *fre* gene does the same function in *S*. Typhimurium by reducing the Fe^+3^ of siderophores to F^+2^ from *fepBDGC* system. The *ggt* is another gene that *S*. Typhimurium uses to cope with iron restriction. The product encoded by *ggt* (γ-glutamyltranspeptidase) is an important enzyme in glutathione metabolism and is required for fitness under iron-restricted conditions. It has been suggested that *ggt* plays a role in Fe-S cluster biosynthesis in *Saccharomyces cerevisiae* [39]. In *Campylobacter jejuni*, *ggt* contributes to the colonization of gut in chicken and humans [40]. *Helicobacter pylori* lacking *ggt* fails to persistently colonize the stomach of mice [41]. The role of glutathione in iron trafficking was previously reported in *S*. Typhimurium and *E*. *coli* [42, 43] but to the best of our knowledge this is the first report on the *ggt* role in *Salmonella* growth in iron restricted conditions. We also identified the genes that import iron from extracellular media. A siderophore gene *fepD* encodes iron enterobactin transporter membrane protein in *E*. *coli* [44]. *fepD* was required for *S*. Typhimurium growth in iron-restricted conditions. In *S*. Typhimurium FepD is a part of FepDGC ABC transporter and is involved in the uptake of siderophore salmochelin [45]. Previously, we showed that *fepD* is important for the bacterium to resist oxidative stress [21]. FepD interacts with TonB and ExbB (**Fig. S4**). *tonB* and exbB were also required for *S*. Typhimurium growth under iron-restricted conditions (**Table 1**). It has been suggested that siderophore complexes depend on TonB to energize the active transport across the membrane via TonB-ExbB-ExbD complex [46]. In *S*. Typhimurium *tonB*-mediated iron uptake is involved in the colonization of this pathogen in the Peyer’s patches and mesenteric lymph nodes of mice [47]. This complex interacts with Suf system via interactions of SodA-NfuA (**Fig. S4**).

#### Sigma E factor

We found *ropE* and *degS* are required for *S*. Typhimurium fitness under iron-restricted conditions. *rpoE* encodes RNA polymerase sigma E factor, while *degS* encodes serine endoprotease. In *E*. *coli*, *rpoE* and *degS* are essential genes; RpoE is an extra-cytoplasmic factor that activates in response to envelope stress. The activation starts by unfolding outer membrane proteins (OMPs) and ends with proteolysis of anti-sigma E factor by DegS to free RpoE and initiate transcription [48, 49]. In *S*. Typhimurium *ropE* and *degS* are not essential genes [24]. RpoE responds to a variety of extra-cytoplasmic stresses in bacteria and the role of this sigma factor has been determined for pathogenesis and virulence; in *Salmonella* the expression of *rpoE* is activated by different types of stressors [50, 51]. Remarkably, our findings demonstrated that Δ*rpoE* mutant is attenuated completely under severe iron-restricted conditions (Dip400). This is reflected in the fact that no sequence reads were detected under Dip 400 (**Supplementary Data set 2**). To confirm the phenotype of these two genes, we grew single deletion mutants of *degS* and *rpoE* in LB supplemented with varying concentrations of Dip. After 24 h growth in a 96-well plate, we did not observe a significant change in growth rate and Max OD_600_ between the mutants and wild-type. Therefore, we used spot dilution assay to confirm Tn-seq results for *degS* and *rpoE* in *S*. Typhimurium. As expected, *S*. Typhimurium *degS* and *rpoE* mutants exhibited growth defects in the presence of Dips as compared to wild-type (**Fig. 3**). Whereas Dip100 did not exhibit the expected phenotype, Δ*degS* and Δ*rpoE* formed smaller colonies at Dip200 as compared to wild-type. At Dip300, both mutants showed a strong phenotype as compared to wild-type. At Dip400, Δ*degS* and Δ*rpoE* did not grow while wild-type did grow slowly. The results of this spot dilution assay confirm that Δ*degS* and Δ*rpoE* are playing important roles in *S*. Typhimurium growth under iron-restricted conditions. The lack of discernable difference in the growth phenotype in broth media, which was captured in the spot dilution assay, indicates the high resolution of our Tn-seq assay in detecting small differences in the mutant fitness within the population of Tn5 mutants. It was previously shown that *S*. Typhimurium lacking *degS* survives very poorly in the macrophages and was slightly attenuated in mice as compared to the wild-type strain, whereas *S*. Typhimurium Δ*rpoE* was attenuated more than 500-fold as compared to *degS* mutant in the mice [50]. In addition, *degS* plays an important role in *S*. Typhimurium survival in elevated temperatures and is required for full virulence [52]. *rpoE* can be activated by acid stress in *S*. Typhimurium and the gene contributes to bacterial survival in the acidified phagosomal vacuole [53]. Microarray analysis indicates that 58% of *S*. Typhimurium genes are affected by *rpoE* and there is a strong connection between SPI-2 and *rpoE* [54]. Also, it has been proposed that *Salmonella* Typhi invasion and intracellular survival are underpinned by *rpoE* via regulation of SPI-1 and SPI-2 [55]. Moreover, it has been suggested that *rpoE* regulates antibiotic resistance in *S*. Typhi through downregulation of the OMP genes and upregulation of the efflux system [56]. Lastly, pertussis toxin (PT) and adenylate cyclase toxin (ACT) are arsenals utilized by *Bordetella pertussis* to kill and modulate host cells and the expression of these two toxins is indirectly modulated by *rpoE* [57]. Altogether, *rpoE* has a broad impact on bacterial fitness and survival in the presence of various host stressors. To the best of our knowledge, this is the first report on the role of *rpoE* in *S*. Typhimurium during iron starvation.

**Figure 3.**
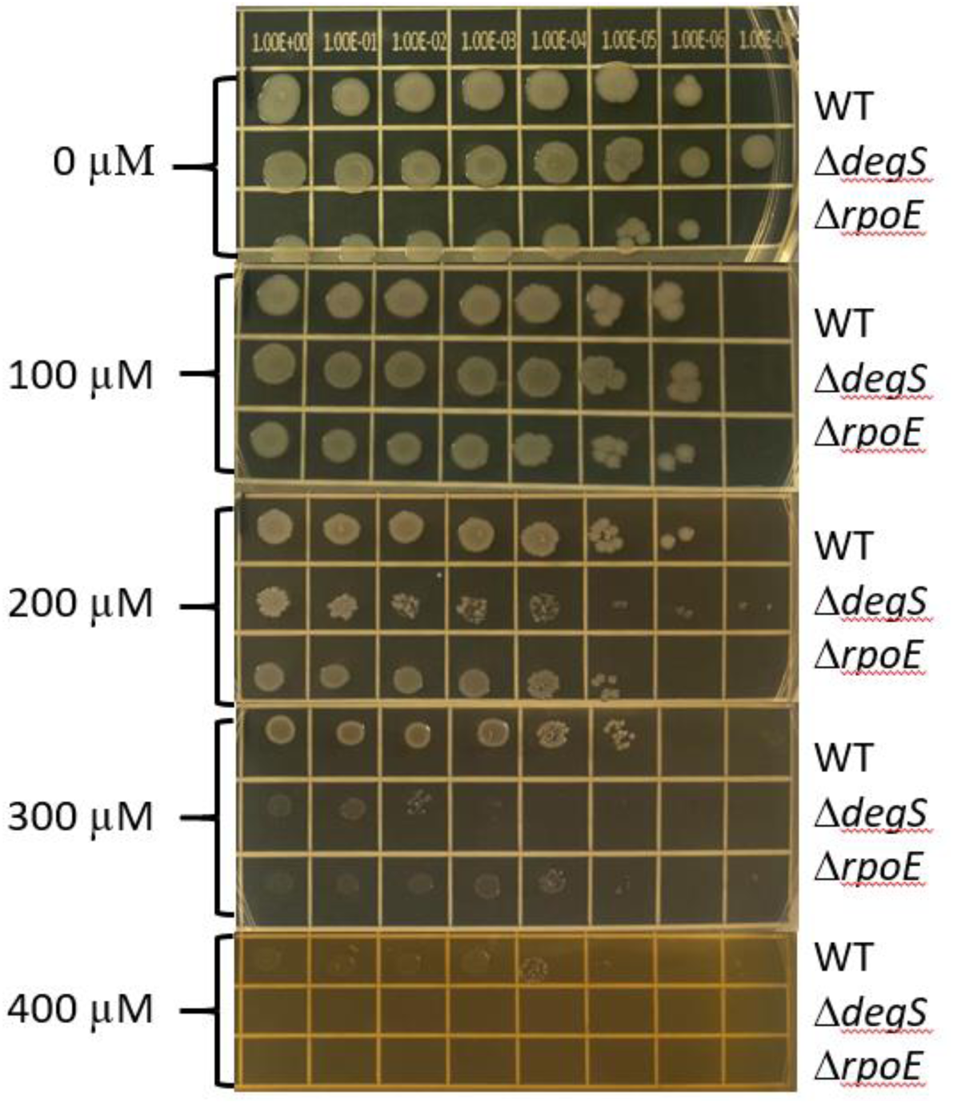
*degS* and *rpoE* are required for the growth *S*. Typhimurium growth under iron-restricted conditions. Spot dilution assay was performed with S. Typhimurium 14028s wild-type, Δ*degS* and Δ*rpoE* mutants. The serial dilutions (10^0^-10^-7^ dilutions) of the overnight cultures of the wild-type, Δ *degS,* and Δ *rpoE*, were spotted on the surface of LB agar plates containing 0 (control), 100, 200, 300, and 400 µM 2,2‵-Dipyridyl (Dip). The plates were incubated at 37 °C and results were recorded after 24 hr.

#### Other miscellaneous pathways

In addition to the genes described in the previous sections, our Tn-seq analysis also identified numerous genes important for fitness under iron-restriction conditions, which are associated with other various biological pathways. These other pathways include (1) oxidative stress, (2) porphyrin biosynthesis, (3) electron transport, (4) gluconeogenesis, (5) glycolysis, (6) osmotic stress, (7) phosphate transport, and (8) Zinc exporter.

Our analysis indicates that *sodA* was required for *S*. Typhimurium growth under iron-restricted conditions. *sodA* gene encodes superoxide dismutase which detoxifies reactive oxygen species. in *E coli* the gene is under control of Ferric Uptake Regulation (Fur) [58]. *S*. Typhimurium Δ*sodA* mutant showed a reduced capacity to invade HeLa cells and form biofilm as well as to resist to chicken serum and reactive oxygen species [59]. It was also previously shown that iron restriction can induce *sodA* expression *in vitro* in *S*. Typhimurium [47]. *hemL* encoding Glutamate-1-semialdehyde 2,1-aminomutase is required for *S*. Typhimurium growth under iron-restricted conditions. The gene is involved in the biosynthesis of 5-aminolevulinic acid from glutamate via the five-carbon pathway [60]. *ydgM* (*rsxB*) is also required for *S*. Typhimurium growth under iron-restricted conditions (Table 1). It has been suggested that the product of *rsxB* plays a major role as the core electron mediators to reduce SoxR (redox-sensitive transcriptional activator), and it is a part of a reductase complex located in the cytoplasmic membrane in *E*. *coli* [61]. The gene *ndh* encoding type II NADH:quinone oxidoreductase (NAHD-II) is required for *S*. Typhimurium to cope with iron-restricted conditions. NAHD-II is a membrane-bound dehydrogenase that plays a central role in respiratory chains in many prokaryotes [62]. Our analysis shows that Ndh interacts with GpmA (**Fig. S4**). Fitness of Δ*gpmA* was reduced under iron-restricted conditions. The gene encodes a phosphoglycerate mutase which is involved in glycolysis and gluconeogenesis. These two genes (*ndh* and *gpmA*) belong to different pathways, and how they interact under iron-deficient environments warrants further investigations. It has been suggested that OsmE is a putative protein regulated by osmotic stress in *E*. *coli* [63]. Our Tn-seq shows that *osmE* is required for *S*. Typhimurium growth under iron-restricted conditions. *pstAB* is also required for *S*. Typhimurium growth under iron-restricted conditions. These genes are parts of ABC transporter *pstSACB* complex which contributes to phosphate import. *pstB* provides energy to the phosphate transporter system via ATP hydrolysis [64]. In *E*. *coli*, *pstSACB* is upregulated in response to phosphate-limited conditions [65]. It is unclear currently how *pstAB* is important for fitness under iron-limited conditions. *zntA* is required for *S*. Typhimurium to grow in iron-restricted conditions. The gene encodes a zinc exporter, which has a role in zinc homeostasis [66]. *zntA* has been implicated in the resistance of *S.* Typhimurium to zinc and copper and is critical for its full virulence [67]. Recent work showed that *S*. Typhimurium ZntA is a part of zinc efflux transporter required to diminish cytotoxic effects of free zinc and to resist nitrosative stress [68]. It was unexpected that our result showed *zntA* is critical for the fitness of this pathogen in iron-deficient media. To validate this finding, *S*. Typhimurium lacking *zntA* was grown in LB media supplemented with 100 and 150 µM Dip (Dip100 and Dip150). The results confirmed our Tn-seq finding and showed that *zntA* is important for the growth of *S*. Typhimurium *in vitro* because the growth rate of the mutants reduced under Dip stress in comparison to the control without Dip (**Fig. 4**). Our Tn-seq results indicates that *zntA* is required for all tested conditions, from the mild to the severe iron restriction (**Table 1**). Regarding the specificity of the used Dip in this study, it has been suggested that Dip cannot chelate zinc, excluding the possibility that the requirement of *zntA* is through depletion of zinc [69].

#### Uncharacterized genes

We also found three previously unknown genes to be important for the growth of *S*. Typhimurium under iron-restricted conditions: *STM14_4330*, *STM14_4612*, *ygjQ*, and *yhfK*. These genes are annotated in UniProt databases as follows. *STM14_4330* putative sugar kinase; *STM14_4612* putative cytochrome c peroxidase; *ygjQ* putative integral membrane protein; *yhfK* putative inner membrane protein.

### Dynamics of the genetic requirements in response to iron restriction levels

In this work, we exposed *S*. Typhimurium Tn5 libraries to different concentrations of iron chelator Dip for comprehensive identification of all genes that contribute to the fitness of this pathogen under iron restriction. For convenience, we categorized the 4 iron-restricted conditions we used into 3 levels: mild (Dip100 and Dip150), moderate (Dip250), and severe iron-restriction (Dip400). Since *S.* Typhimurium faces iron limitation at different levels of severity during infection in the host, we reason that our strategy for Tn-seq selection under a wide range of iron restriction can comprehensively capture the genes that are important in coping with the iron stressor at various stages or niches of *Salmonella* infection in the host. Interestingly, we found that not all 28 conditionally essential genes required for fitness under iron restriction were required under all 4 levels of iron-restricted conditions (**Table 1**). The 28 genes were clustered into different groups according to the patterns of the mutant fitness values in response to different levels of iron restriction (**Fig. S5 and Fig. S6**). The genes of Suf system, *sufABCDSE*, which were required under moderate and severe conditions (Dip250 and Dip400), were not required for mild iron restriction conditions (Dip100 and Dip150). The fitness of these mutants was reduced more when iron-restriction severity elevated: the average fitness of these mutants was −1.8 and −2.6 for Dip250 and Dip400, respectively. This suggests that *S*. Typhimurium uses Suf system to survive in moderate and severe iron restricted niches., The fitness of the mutant in *iscR* encoding *sufABCDSE* regulator was reduced in Dip250 and Dip400, −2.75. However, *S*. Typhimurium Δ*iscR* grew better in mild iron-restricted conditions (Dip100 and Dip150) with the fitness of Δ*iscR* +3.09. For the *nfuA*, the gene was required only under moderate iron-restricted conditions.

Interestingly, the fitness of the siderophore gene *fepD* was −2.91 in Dip100 and reduced to −4.27 in Dip150 and −3.56 in Dip250, while not required in D400 for *S*. Typhimurium growth. This suggests that *fepD* is critical for bacterial growth under mild and moderate iron-restricted conditions but not under severe iron restriction. Δ*tonB* exhibited similar phenotype as *fepD* with fitness of −6.43, −4.32 and +1.44 for Dip150, Dip250 and Dip400, respectively. Δ*exbB* also behaved similarly. Unexpectedly, the sigma E factor had the highest reduced fitness in moderate iron-restricted conditions (the fitness −5.965) and it reduced further in severe iron-restricted conditions (−7.24). Also, the fitness of Δ*degS*, the mutant without functional anti-sigma E factor, was −2.605 for Dip250, and Δ*degS* recorded the lowest fitness of −8.35 in Dip400. This demonstrates that *rpoE* and *degS* are increasingly required for *S*. Typhimurium growth under both iron-restricted conditions in a manner dependent on the concentration of Dip. The 6 genes, *sodA*, *gpmA*, *ydgM*, *yhfK*, *STM14_0026*, and *STM14_4612*, were only required in moderate iron-restricted conditions for *S*. Typhimurium growth. Whereas *ggt*, *hemL*, *ndh*, *osmE*, *STM14_4330*, and *ygjQ* were only required in severe iron-restricted conditions. The fitness of *pstB* in moderate iron-restricted conditions was −1.395 and in severe iron-restriction, it became −2.04 while *pstA* was required only in severe iron restriction conditions for *S*. Typhimurium growth. Finally, *zntA* was significantly required in Dip150 and Dip400, with the fitness values of −1.39 and −1.03. Altogether, these findings demonstrate the requirements of the genes identified in this study for fitness are dependent on the severity of iron restriction and suggests that certain genes might be required for fitness under a specific range of iron restriction.

### Genes that when deleted increase fitness under iron-restricted conditions

In this study, our main goal was to identify the genes that when deleted reduce the fitness under iron-restricted conditions. However, we also identified 30 genes that when the deleted result in increased fitness: that is, their mutants grow better under iron-restricted conditions. These phenotypes were mainly observed in moderate (Dip250) and severe iron-restricted conditions (Dip400)(**Supplementary Data set 1** and **Table 2**). We briefly categorize them based on their biological functions and highlight a few of these genes. First, the genes involved in nucleotide biosynthesis and metabolism: the fitness of the subunits of DNA polymerase V, *umuD* was increased under iron-restricted conditions. DNA polymerase V incorporates 8-oxo-guanine into DNA during replication, it has been shown that *E*. *coli* lacking *umuD* confers resistance to the antibiotics and can grow in presence of the antibiotics [70]. In addition, the fitness of Δ*guaB* and Δ*purA* also increased. These two genes catalyze the first step in the *de novo* synthesis of guanine and adenine from inosine 5’-phosphate (IMP). Second, the genes involved in TCA cycle: we found that the mutant fitness of *acnB*, *icdA*, and *sdhD* genes, which encode TCA cycle enzymes, increased under iron-restricted conditions; *S*. Typhimurium lacking one of these genes can grow better under iron-restricted conditions, whereas the fitness of other mutants in TCA cycle did not change. A similar occurrence was observed when *E*. *coli* was exposed to lethal doses of antibiotics. The survival of the deletion mutants in *acnB* or *icdA* increased against bactericidal antibiotics [11]. Third, the genes involved in carbohydrate metabolism: *S*. Typhimurium strains with a deletion in *yaeD*, *eda*, or *STM14_2709* can grow better under iron-restricted conditions. *eda* encodes Entner-Doudoroff aldolase which has a central role in sugar acid metabolism and detoxification of metabolites in *E*. *coli* [71]. Finally, the genes involved in various pathways or functions, including lipopolysaccharide biosynthesis (*rfbB*, *rfaC*, *rfbH,* and *rfbI*), integral component of membrane (*eyeiB*, *ychH*, *STM14_0726*), membrane transporters (*sapA*, *ompW*, *smvA*), membrane-localized protease (*htpX*) and outer membrane protein assembly factor (*nlpB*). This suggests that mutations in any of the twelve genes associated with the membrane-related functions led *S*. Typhimurium to grow better under iron-restricted conditions. Among all those genes, only three genes (*acnB*, *rfbI*, and *STM14_0726*) possibly bind to Fe-S. Currently, we are lacking a scientific explanation of how the deletions in these genes caused the bacteria to grow better under iron-restriction stress.

### Conclusions

In this work, we characterized the genome of *S*. Typhimurium at a system-wide level to identify the genes required for growth under the stress of iron restrictions. We also demonstrated the requirements of these genes for fitness alter according to the severity of the iron restriction. We validated the phenotypes only for 4 single deletion mutants for *sufS*, *zntA*, *degs*, and *rpoE* genes. However, our previous works indicated that 84% (50 mutants tested) of the genes identified by our Tn-seq method and bioinformatic pipeline displayed the expected phenotypes. We plan to move with these *in vitro* findings for further evaluation using macrophage cell lines and mice. The results of this study can be exploited for the development of effective therapeutic strategies and it can expand our knowledge about how *Salmonella* survives in iron-restricted environments.

## Materials and Methods

### Growth response of *S*. Typhimurium to different concentrations of 2,2‵-Dipyridyl (Dip)

A single colony of *Salmonella* enterica serovar Typhimurium ATCC14028s was inoculated into a 2 ml LB (Luria-Bertani) broth medium in a 5 ml tube and incubated overnight in dark for 16 h. The single colony was taken always from an agar plate with the colonies not older than 10 days. The next day, freshly prepared LB broth media supplemented with different concentrations of Dip were inoculated with the *S*. Typhimurium 14028s culture at 1:200 dilutions. Dip was dissolved in ethanol and then diluted in autoclaved MQ-H_2_O before adding it to LB broth media. The cultures (200 μl/well) were immediately added into a 96-well microplate, incubated in a Tecan Infinite 200 microplate reader at 37°C with shaking amplitude of 1.5mm and shaking duration of 5 s. OD_600_ of the cultures were measured every 10 min. After 18 h incubation, the data were collected and run on GrowthRates script to calculate growth rate, and maximum OD_600_ [72].

### Preparation of *Salmonella* Typhimurium Tn5 mutant libraries

Tn5 mutant libraries were constructed as previously described [21, 22]. Briefly, a spontaneous mutant strain of *S*. Typhimurium ATCC 14028s, which is resistant to nalidixic acid (NA^R^), was mutagenized by conjugation utilizing *Escherichia coli* SM10 λ*pir* harboring a suicide pBAM1 transposon-delivery plasmid vector (Amp^R^) as the donor cell [73]. One ml of overnight growth cultures from the donor and recipient bacteria each were washed with 10 mM MgSO_4,_ mixed, and concentrated on the nitrocellulose filter. The filter was placed on an LB agar plate, which was then incubated for 5 h at 37°C. After the conjugation, the cells were washed with the MgSO_4_ and plated on LB agar plates containing NA and Km. The plates were grown at 37°C for 24 h in dark. Finally, colonies were scrapped, added into LB broth containing 7% DMSO, and stored at −80°C in aliquots.

### Tn-seq selection conditions of iron restrictions

The aliquot of the Tn5 mutant library was thawed at room temperature on ice and then diluted in LB broth. The library was incubated at 37°C with shaking at 225 rpm for an hour and then washed twice with PBS. This allows the mutants to reactivate and get rid of DMSO residues. The library was inoculated to 20 ml LB broth in a 300 ml flask and LB broth supplemented with 100, 150, 250, or 400 μM Dip (Dip100, Dip150, Dip250, and Dip400, respectively), and LB broth without Dip was used as a control. The inoculum of each library represented approximately 10 cells for each mutant. Initially, we prepared a library (Library-A) then another library (Library-B), and both were combined to form Library-AB. Library-A was used for selection conditions Dip100, Dip150, and LB-II (Control), while Library-AB was used for the conditions of Dip250-I, Dip250-II, Dip400, and LB-III (Control). The cultures were incubated at 37°C with shaking at 225 rpm in a dark and humidity-controlled incubator till they reached the mid-log phase. Then, the cultures were pelleted and stored at −20°C.

### Tn-seq library generation for HiSeq sequencing

Previously we developed a robust method for Tn-seq library preparation, and therefore the Tn-seq amplicon libraries for HiSeq sequencing were prepared according to the protocol [21, 22]. Bacterial genomic DNA was extracted for each condition utilizing DNeasy Blood & Tissue kit (Qiagen). Qubit dsDNA RB Assay kit was used to quantify extracted genomic DNA (Invitrogen). Genomic DNA was digested with PvuII-HF (New England Biolabs) and purified with DNA Clean & Concentrator-5 kit (Zymo Research), this step is critical for removing the reads from pseudo Tn5 mutants. Next, a linear PCR extension was conducted utilizing Tn5-DPO primer (**Table S4**). The PCR reaction was conducted in a 50 μl contained Go Taq Colorless Master Mix (Promega), 20 μM Tn5-DPO primer, 100 ng gDNA, and MQ-H_2_O. The PCR cycles were 95°C for 2 min, followed by 50 cycles at 95°C for 30 sec, 62°C for 45 sec, and 72°C for 10 sec. The amplicon was purified with DNA Clean & Concentrator-5 kit. The C-tailing reaction was performed by using terminal transferase (TdT) buffer (New England Biolabs), CoCl_2_, dCTP, ddCTP, TdT, and the purified linear PCR product. The mixture was incubated at 37°C for 1 h, followed by inactivation of TdT by incubation at 70°C for 10min. The C-tailed product was purified using the DNA Clean & Concentrator-5 kit. Next, the exponential PCR was conducted utilizing P5-BRX-TN5-MEO and P7-16G primers (**Table S4**). The PCR reaction was conducted in a 50 μl using Go Taq Green Master Mix, P5-BRX-TN5-MEO and P7-16G primers, the C-tailed genomic junctions, and MQ-H_2_O; the PCR cycles were 95°C for 2 min, followed by 30 cycles of 95°C for 30 sec, 60°C for 30 sec, and 72°C for 20 sec, and the final extension at 72°C for 5 min. Next, the PCR products were run on an agarose gel and the DNA fragments size 325 – 625 bp were extracted using Zymoclean Gel DNA Recovery kit (Zymo Research). The DNA libraries were quantified utilizing Qubit dsDNA RB Assay kit. The libraries were pooled and sequenced on a flow cell of HiSeq 3000 Illumina utilizing single-end read and 151 cycles at the Center for Genome Research & Biocomputing in Oregon State University.

### Tn-seq data analysis

The output data of the Hi-Seq sequencer were downloaded onto the High-Performance Computing Center (AHPCC) at the University of Arkansas. Since samples were multiplexed before sequencing, a custom Python script was used for de-multiplexing them. The script searched for the six-nucleotide barcode of each library with no mismatch allowed. Tn-Seq Pre-Processor (TPP) tool was utilized to extract transposon genomic junctions [74]. We modified the script of the TPP to process our sequences. In a fixed sequence window, TPP searched for the 19-nucleotide Tn5 inverted repeat (IR) and identified the five nucleotides GACAG at the end of the IR sequence. The genomic junctions of *S*. Typhimurium that start immediately after GACAG were extracted and the C-tails were trimmed. The genomic junction sequences of less than 20 nucleotides were excluded and the remaining junction sequences were mapped to *Salmonella* enterica serovar Typhimurium 14028s genome utilizing BWA-0.7.12 [75]. Essentiality Indices (EI) were calculated using Tn-seq Explorer [23].

### Identification of conditionally essential genes required for growth under iron-restricted conditions

The conditionally essential genes were identified utilizing TRANSIT tool [74]. The resampling algorithm was utilized for the analyses in TRANSIT. The LB-II and LB-III were the inputs (controls), while Dip100, Dip150, Dip250-I, Dip250-II, and Dip400 were the outputs (experiments). Trimmed Total Reads (TTR) were used as a normalization method and 10,000 samples were used for the analysis. The insertions in 5% of N-terminal and 10% of C-terminal of ORFs were excluded. Genes were considered conditionally essential when *p* values were < 0.05.

#### Phenotypic analyses of the single deletion mutants

For assessment of the growth phenotypes of the mutants in LB broth, the overnight cultures of the wild-type *S*. Typhimurium, Δ*sufS*, and Δ*zntA* were inoculated into LB broth containing 0, 100, or 150 µM Dip with the inoculum diluted at 1:200. Then, 200 µl of cultures were directly added into 96-well microplates and incubated in Tecan infinite 200. The incubation time was 18 h at 37°C. The OD_600_ data was collected every 10 min and used to calculate the growth rate with GrowthRates script [72] and to obtain the maximum OD_600_. For the spot dilution assay, the overnight cultures of the wild-type, Δ *degS*, and Δ *rpoE* were serially diluted from 10^0^ to 10^-7^ in a 96-well plate. LB agar plates were prepared two days earlier to contain 0, 100, 200, 300, and 400 µM Dip. Five µl of serially diluted cultures were spotted on the agar plates and let dry completely at room temperature. The plates were incubated at 37 °C and results were recorded after 24 h. All single deletion mutants were obtained through the NIH Biodefense and Emerging Infections Research Resources Repository, NIAID, NIH: *Salmonella enterica* subsp. *enterica*, Strain 14028s (Serovar Typhimurium) Catalog No. NR-42850, NR-42853.

## Availability of data

The bam file of all seven conditions is available on NCBI SRA under BioProject number PRJNA397775.

## Author Contributions

Conceived and designed the experiments: SK, YK, Performed the experiments, analyzed the data, wrote the manuscript: SK, TJ, Revised the manuscript: SK, YK

## Supplementary Data sets

**Supplementary Dada set 1.** The genes in *S*. Typhimurium 14028s that are identified in this study to be implicated in the growth under iron-restricted conditions. Sheet 1 (“Reduced fitness”) shows the genes that are required for the fitness of the wild-type strain. Sheet 2 (“Increased fitness”) shows the genes that lead to increased fitness of the respective mutants. Iron-restricted conditions were achieved by supplemented LB broth with 2,2‵-Dipyridyl (Dip) at the concentrations of 100, 150, 250, and 400 µM. Cultures were grown at 37°C till the mid-log phase.

**Supplementary Data set 2.** The summary of the Tn-seq data analysis for all genes in the genome of *S.* Typhimurium 14028s for each iron restriction condition. For Tn-seq selection, Tn5 libraries of *S*. Typhimurium 14028s were selected in LB broth supplemented with 2,2‵-Dipyridyl (Dip) at the concentrations of 100, 150, 250, and 400 µM until the mid-log phase.

## Supplementary Materials

**Table S1.** Effect of different concentrations of 2,2‵-Dipyridyl (Dip) on the growth of the wild-type S. Typhimurium 14028s.

| Condition | N | Growth Rate (GR) | Max OD<sub>600</sub> | GR % Reduction | Max OD<sub>600</sub> % Reduction |
| --- | --- | --- | --- | --- | --- |
| **LB** | 7 | 0.0240 | 0.938 | 0.00 | 0.00 |
| **Dip100** | 8 | 0.0227 | 0.897 | 5.37 | 4.39 |
| **Dip150** | 8 | 0.0220 | 0.834 | 8.26 | 11.00 |
| **Dip250** | 8 | 0.0201 | 0.615 | 16.30 | 34.40 |
| **Dip400** | 7 | 0.0176 | 0.488 | 26.40 | 48.00 |

**Table S2.** The time required for Tn-seq selection cultures to reach the mid-log phase.

|  | **Time to reach mid-log phase** | **OD<sub>600</sub>** |
| --- | --- | --- |
| **Library-A (325,000 mutants)** |  |  |
| **LB-II** | 5 hr 0 min | 2.63 |
| **Dip100** | 5 hr 35 min | 2.61 |
| **Dip150** | 6 hr 5 min | 2.57 |
| **Library-AB (650,000 mutants)** |  |  |
| **LB-III** | 5 hr 30 min | 2.57 |
| **Dip250-I** | 10 hr 0 min | 2.45 |
| **Dip250-II** | 10 hr 0 min | 2.46 |
| **Dip400** | 12 hr 0 min | 1.84 |

**Table S3.** The summary of the HiSeq sequencing reads.

|  | **Reads with**<br>**Tn5** | **Extracted**<br>**reads** | **Mapped**<br>**Reads** | **Unique Insertions** |
| --- | --- | --- | --- | --- |
| **LB-II** | 38,808,640 | 31,728,005 | 25,223,444 | 125,449 |
| **Dip100** | 25,788,698 | 21,034,947 | 16,991,894 | 117,474 |
| **Dip150** | 36,677,408 | 29,905,496 | 24,364,738 | 121,132 |
| **LB-III** | 57,779,778 | 47,575,248 | 39,248,662 | 193,728 |
| **Dip250-I** | 29,832,849 | 25,082,465 | 21,096,630 | 179,562 |
| **Dip250-II** | 35,439,669 | 28,119,351 | 23,104,233 | 181,534 |
| **Dip400** | 30,028,187 | 26,382,625 | 23,135,546 | 169,666 |
| **Total** | 254,355,229 | 209,828,137 | 173,165,147 | 1,088,545 |

**Table S4.**
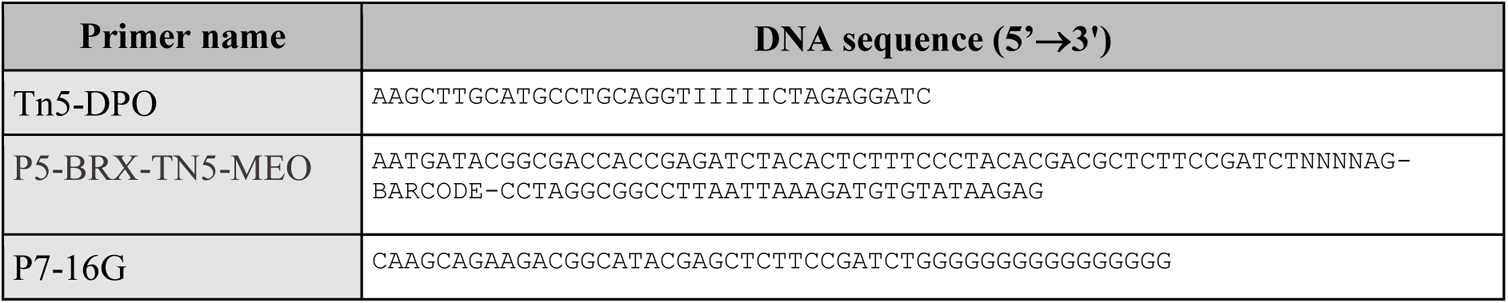
The primers used for Tn-seq library preparation.

**Figure S1.**
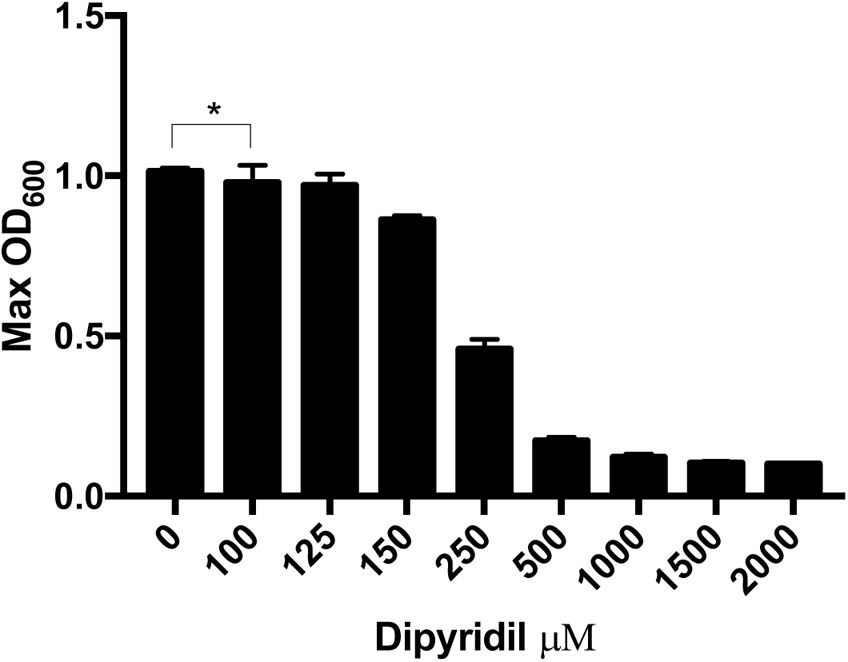
*S*. Typhimurium growth response to 2,2‵-Dipyridyl (Dip). *S*. Typhimurium 14028s wild-type was grown in LB broth supplemented with a 2,2‵-Dipyridyl (0, 100, 125, 150, 250, 500, 1000, 1500, or 2000 µM). The cultures were in a 96-well plate were incubated at 37°C in a Tecan Infinite 200 microplate reader. Maximum OD_600_ (Max OD_600_) was recorded after 18 hr incubation. Statistical significance was determined by unpaired two-tailed *t* test, \**P* < 0.05.

**Figure S2.**
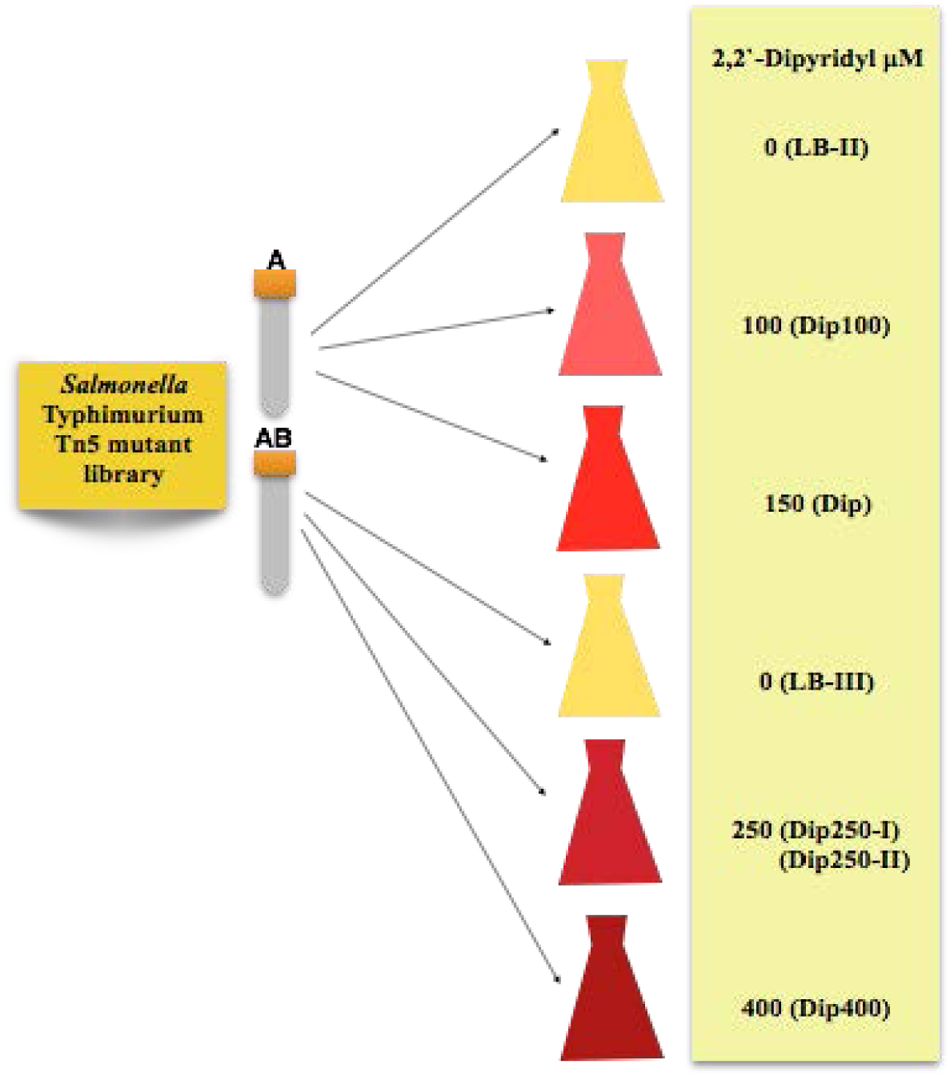
Schematic representation of Tn-seq study design using different levels of iron restriction conditions. Tn5 mutant libraries of *S*. Typhimurium (Library-A and Library-AB) were used to inoculate LB broth supplemented with 2,2‵-Dipyridyl (Dip) at the concentrations of 100, 150, 250, and 400 µM. The controls are free of Dip. The cultures were grown until the mid-log phase. LB-II and LB-III were used as Inputs (controls) and the rest of the cultures were as Outputs for the generation and comparative analysis of the Tn-seq profiles.

**Figure S3.**
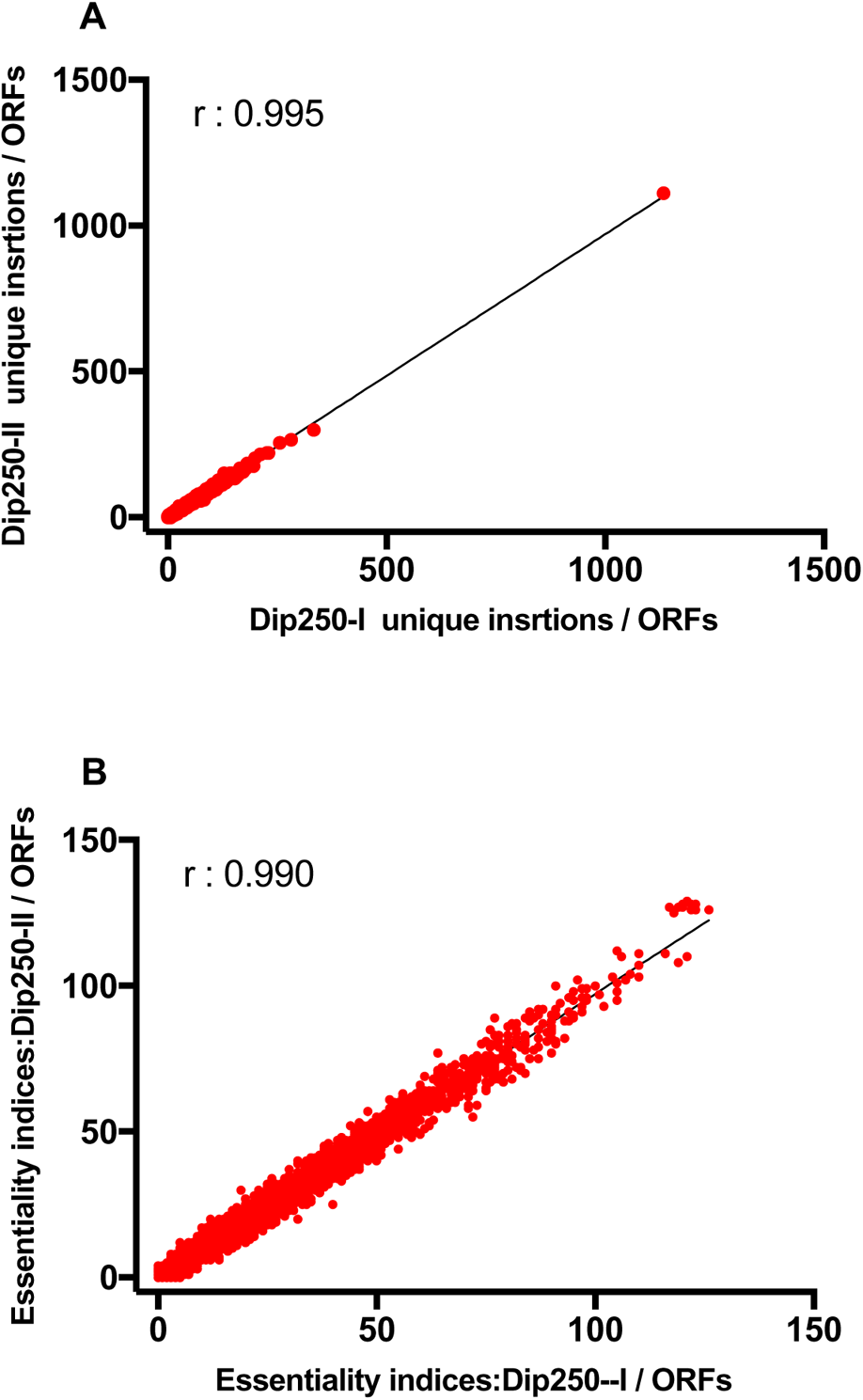
Reproducibility of the Tn-seq. Pearson correlation of the Tn-seq profiles between the two biological replicates obtained from Dip250, Dip250-I, and Dip250-II. (A) correlation between the unique insertions of Dip250-I and Dip250-II for each open reading frame (ORF). (B) correlation between the essentiality indices of Dip250-I and Dip250-II for each open reading frame (ORF).

**Figure S4.**
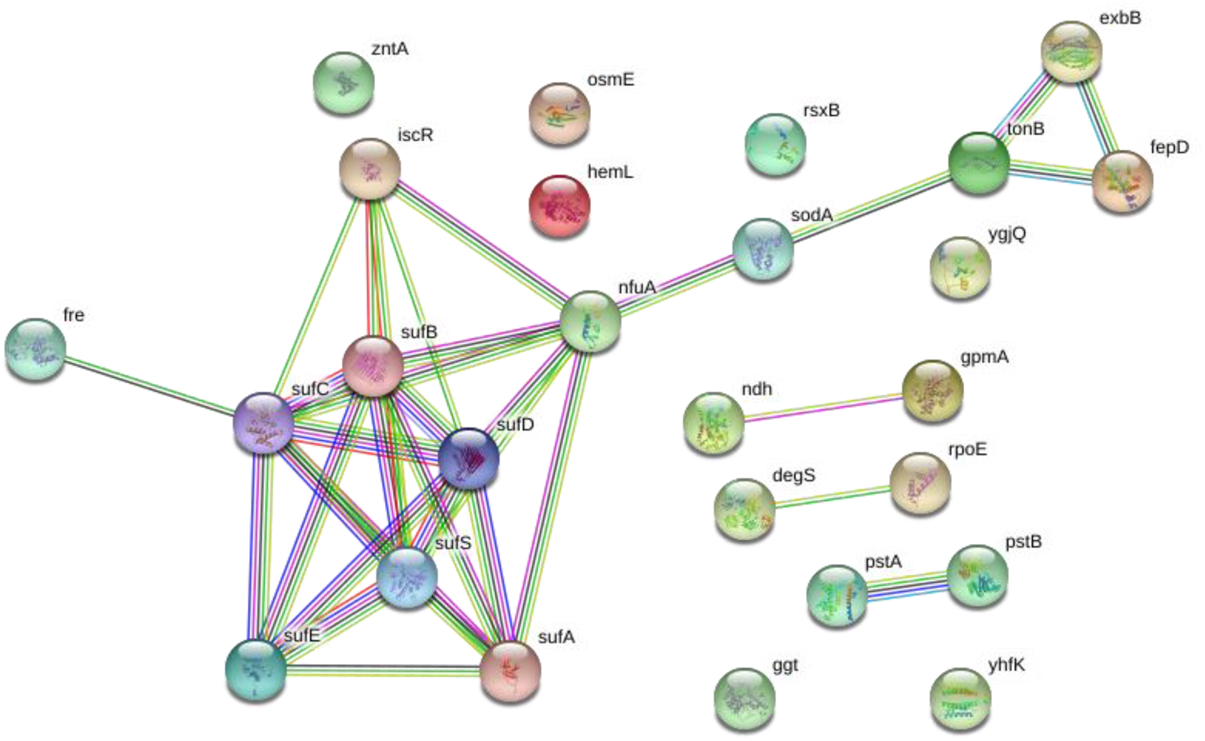
Protein-protein interaction network of the genes required for the growth of *S*. Typhimurium 14028s under iron restriction conditions. The list of the genes was as the input into String protein-protein interaction database using default options. The interactions indicate that Fe-S cluster proteins interact with siderophore proteins via NfuA and SodA.

**Figure S5.**
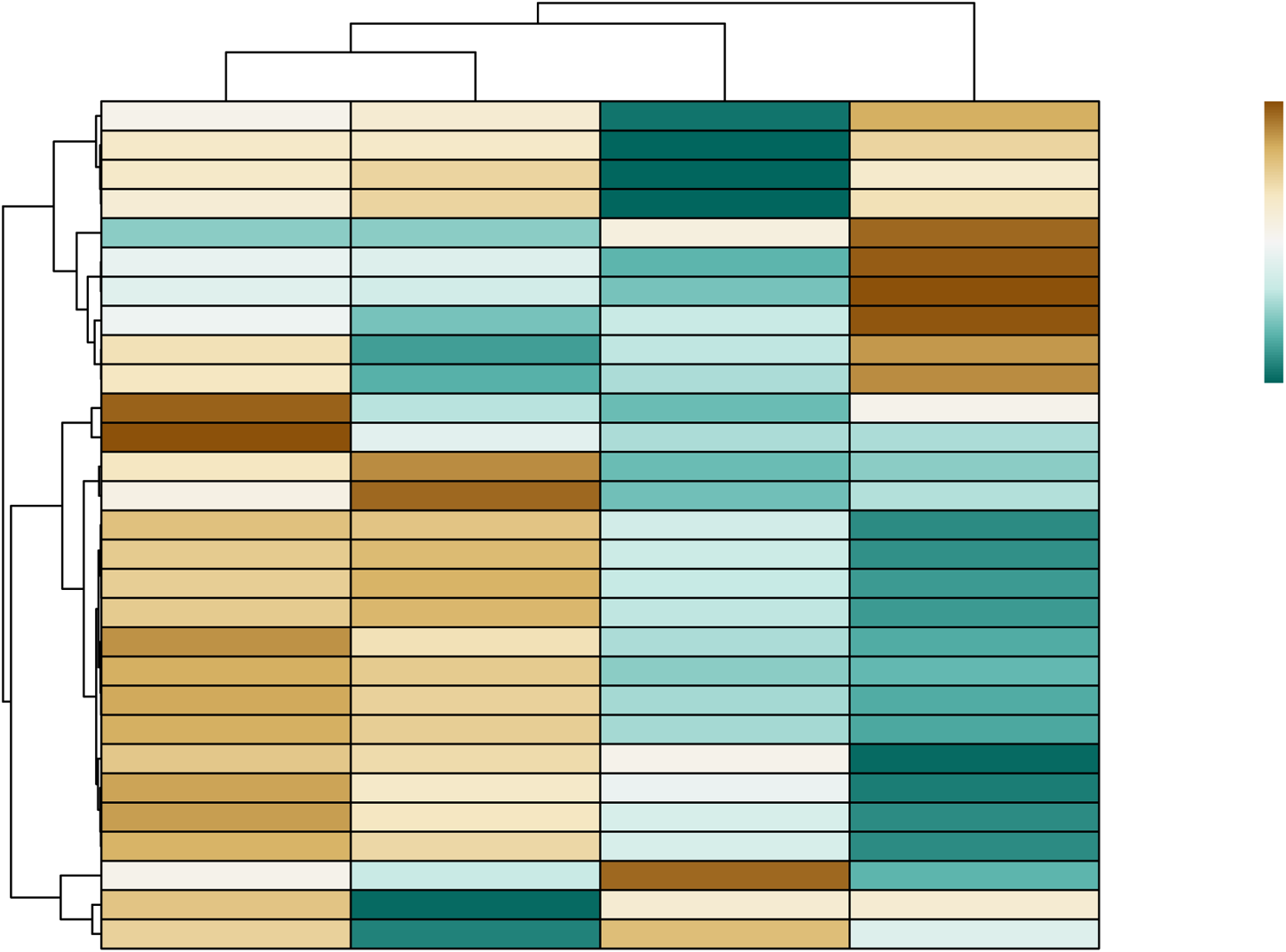
Clustering analysis of the genes in *S.* Typhimurium 14028s identified in this study as conditionally essential for the growth under 4 different levels of iron restriction condition. The 28 conditionally essential genes were clustered according to the changes in the fitness value of the respective mutants in response to 4 different levels (100, 150, 250, and 400 µM Dip) of iron restriction. Unit variance scaling was applied to the fitness values. Both rows and columns are clustered using correlation distance and average linkage. The clustering diagram was generated using ClustVis (biit.cs.ut.ee/clustvis).

**Figure S6.**
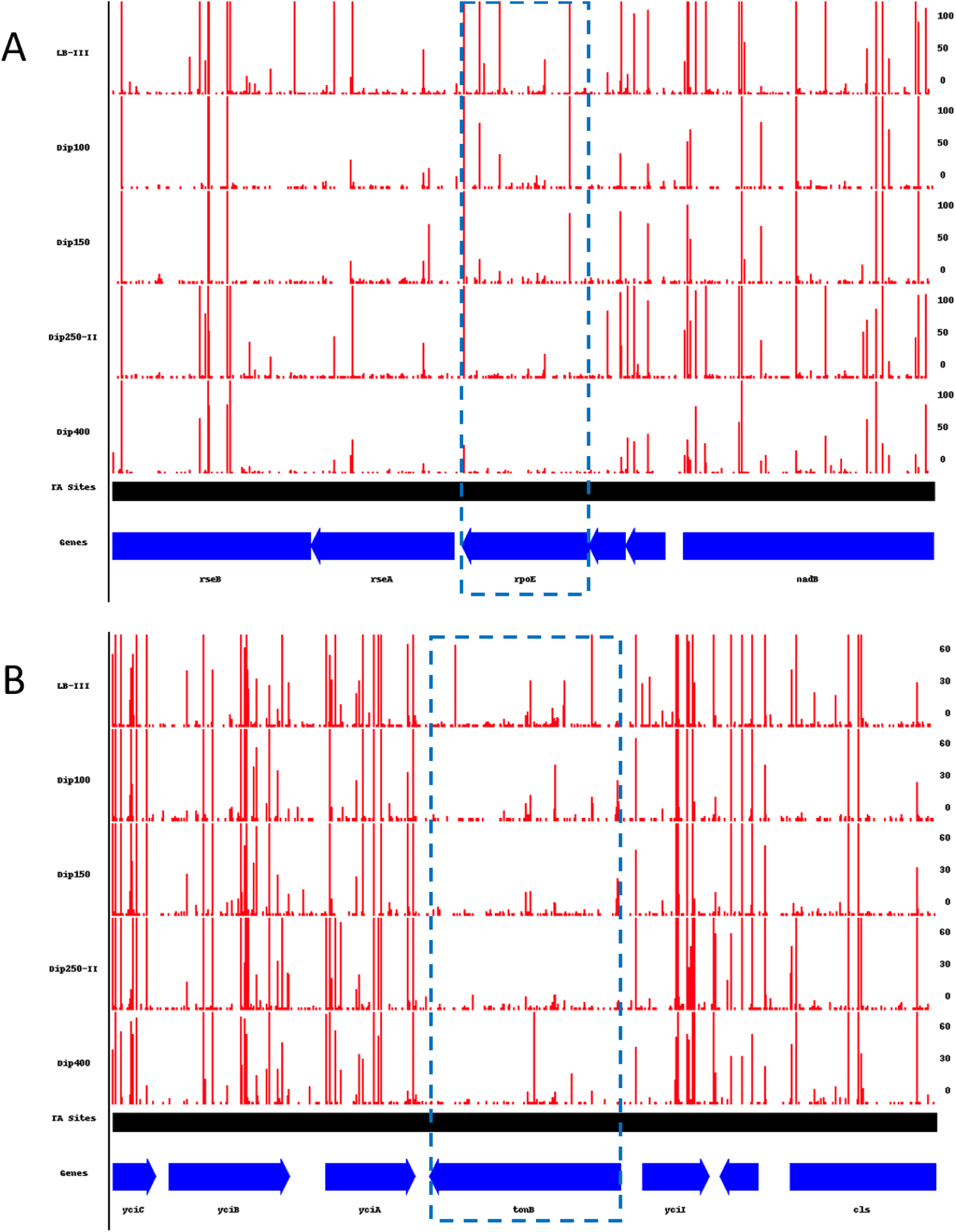
Comparison of the Tn-seq profiles for (A) *rpoE* and (B) *tonB* and their surrounding genes across different levels of iron restriction (100, 150, 250, and 400 µM Dip)

